# Inception Capsule Network for Retinal Blood Vessel Segmentation and Centerline Extraction

**DOI:** 10.1101/815555

**Authors:** C. Kromm, K. Rohr

## Abstract

Automatic segmentation and centerline extraction of retinal blood vessels from fundus image data is crucial for early detection of retinal diseases. We have developed a novel deep learning method for segmentation and centerline extraction of retinal blood vessels which is based on the Capsule network in combination with the Inception architecture. Compared to state-of-the-art deep convolutional neural networks, our method has much fewer parameters due to its shallow architecture and generalizes well without using data augmentation. We performed a quantitative evaluation using the DRIVE dataset for both vessel segmentation and centerline extraction. Our method achieved state-of-the-art performance for vessel segmentation and outperformed existing methods for centerline extraction.

## 1. INTRODUCTION

The status of retinal vessels is an important indicator for many ophthalmological diseases, and accurate vessel segmentation is crucial. Since manual segmentation is time-consuming and error-prone, automatic methods are required [1]. However, segmentation and centerline extraction of retinal blood vessels are challenging tasks due to thin vessel branches and overlapping or crossing vessels [2].

Existing state-of-the-art methods for retinal vessel segmentation use mostly deep convolutional neural networks (CNNs). These methods outperform classical approaches based on handcrafted features and exceed human-level performance (e.g., for image classification [3]). Most methods use a CNN architecture and image patches to classify the central pixel as either vessel or background (e.g., Melinščak et al. [4]). Fu et al. [5] reformulated the problem of vessel segmentation as a boundary detection task using a CNN in combination with a conditional random field (CRF) to classify image patches. Tetteh et al. [6] used the Inception architecture without pooling layers to classify each pixel in an image patch and extracted vessel centerlines. CNN autoencoder networks also classify each pixel of an image patch (e.g., [7], [8], [9]). The recent approach by Yan et al. [9] takes into account the thickness of vessels by jointly adapting segment-level and pixel-wise loss functions. It has also been shown that appropriate pre-processing, data augmentation, and ensembles of multiple networks can significantly improve the performance, but increase the training time [10]. While deep CNNs have proven to be very good feature extractors, they suffer from several disadvantages such as lack of robustness against affine transformations, and loss of relevant information about spatial relations of pixels due to max-pooling layers. An alternative to CNNs was recently introduced by Sabour et al. [11]. There, the concept of a Capsule was proposed, which is a group of neurons whose output is not just a scalar but a vector which represents specific characteristics of an object. Unlike CNNs, Capsule networks do not employ max-pooling layers but instead use an iterative routing-by-agreement algorithm that determines the coupling of Capsules between consecutive layers. It was shown that a shallow Capsule network outperforms a deep CNN on the MNIST dataset (handwritten digits). Capsule networks have fewer parameters and also require less training data by learning viewpoint invariant feature representations as shown by Hinton et al. [12].

In this contribution, we propose a new deep learning method for segmentation and centerline extraction of retinal blood vessels. Our method combines the recently proposed Capsule network [11] with an Inception architecture [13]. Compared to previous methods on retinal vessel segmentation, the introduced Inception Capsule network is based on a shallow architecture that has fewer parameters and does not require data augmentation to achieve good performance. To our knowledge, it is the first time that a Capsule network is used for retinal vessel segmentation and centerline extraction. We evaluated the performance of our method using the DRIVE dataset [14]. It turned out that our Inception Capsule network yields state-of-the-art results for vessel segmentation, and improves the results for centerline extraction.

## 2. METHODS

Our deep learning method for segmentation and centerline extraction of retinal blood vessels is based on image patches and combines the Capsule network with the Inception architecture. Below, we first describe the architecture of Capsule networks. Then, we detail our newly proposed model and the training procedure.

### 2.1. Capsule Networks

As introduced in [11], a Capsule is a group of neurons whose output is not just a scalar but a vector that represents properties of an object (or part of an object) in an image, for example, color, velocity, or deformation strength. The length of the vector represents the probability that an object is present in a given image or not. A Capsule network (CapsNet) consists of a convolutional layer followed by a composition of multiple Capsule layers. The first Capsule layer is called PrimaryCapsule. Each Capsule layer consists of several stacked Capsules that receive input from lower level Capsules. Instead of using a standard non-linear activation function like ReLU, CapsNet uses a squashing function that squashes the output vector to a length between 0 and 1. Given the total input *s*_*j*_ to Capsule *j* from the previous layer *l*, the vector output *v*_*j*_ of Capsule *j* in layer *l* + 1 after applying the squashing function is defined by

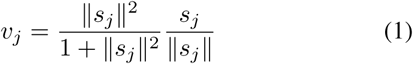

To determine the input to a Capsule in layer *l* + 1, the vector outputs of the Capsules in the previous layer *l* are combined by a weighted sum using coupling coefficients:

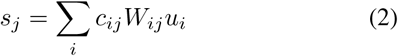

where *u*_*i*_ is the prediction vector from Capsule *i* in layer *l, W*_*ij*_ is the weight matrix that is learned, and *c*_*ij*_ are the coupling coefficients. An iterative routing-by-agreement algorithm [11] determines the coupling coefficients between consecutive Capsule layers. If the scalar product between the output vector of a lower level Capsule and the prediction vector of the next higher level Capsule is large, the coupling coefficient between these Capsules is increased. The sum over all *c*_*ij*_ is 1. To improve the performance of Capsules, Sabour et al. [11] added a subnetwork that reconstructs the input image given the true label.

### 2.2. Inception Capsule Network

Our deep learning method extends the Capsule network architecture and adapts it for retinal blood vessel segmentation and centerline extraction. The method combines the Capsule network architecture [11] with an Inception network [13]. In contrast to the standard Capsule network that uses a convolutional layer before the PrimaryCapsule layer, our Capsule Inception network employs an Inception layer. In addition, compared to the PrimaryCapsule layer in [11], our Inception PrimaryCapsule layer incorporates three Capsule layers with different kernel sizes. Fig. 1 shows the PrimaryCapsule layer as used in our method. The output of each PrimaryCapsule layer is routed to the next Capsule layer via iterative routing [11]. Fig. 2 depicts the Inception PrimaryCapsule layer of our method consisting of three standard PrimaryCapsule layers with 1 × 1, 3 × 3, and 5 × 5 kernel sizes.

**Fig. 1:**
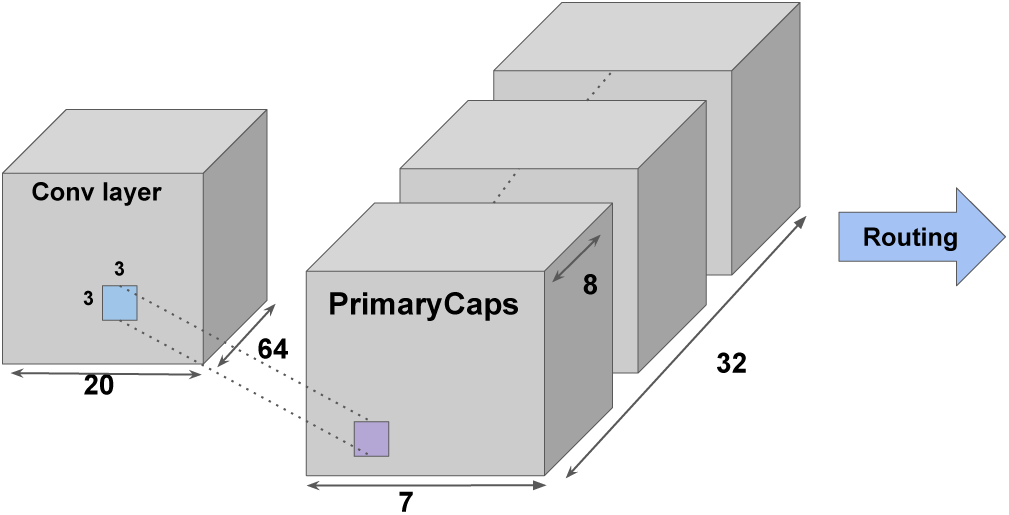
Example of a PrimaryCapsule layer with a 3 × 3 kernel receiving input from a convolutional layer and routing it to the next Capsule layer.

**Fig. 2:**
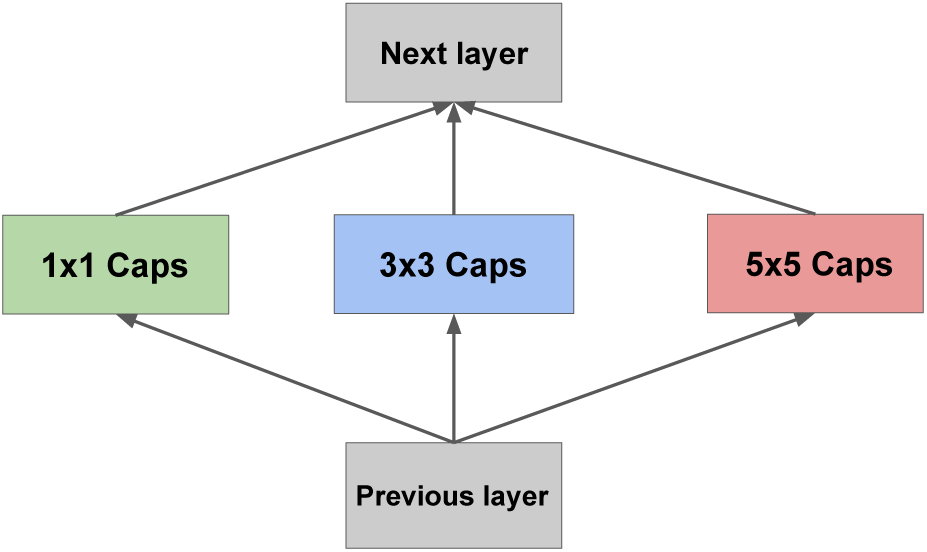
Proposed Inception PrimaryCapsule layer.

The architecture of the proposed Inception Capsule network is shown in Fig. 3. For the first convolutional layer we used an Inception V4 layer [15], which receives the image patches as input. This layer contains 32 filters for each kernel size *k* × *k* with *k* = 1, 3, and 5 with ReLU non-linearity and an identity mapping like in a ResNet block. The output of the Inception V4 layer serves as input for the Inception PrimaryCapsule layer. Each Capsule layer in the Inception PrimaryCapsule layer consists of a convolutional layer with 32 channels, where each channel is a Capsule consisting of 8 convolutional layers (defined by the kernel sizes *k* = 1, 3, 5), see Fig. 2. After the Inception PrimaryCapsule layer, a final Capsule layer (VesselCaps) is used with 12-dimensional Capsules. Using Inception PrimaryCapsules increases the number of parameters (10.3 million) compared to CapsNet [11] (6.8 million), however, compared to existing methods for retinal vessel segmentation (e.g., the model in [5] involves 32.5 million parameters) the number of parameters is much lower, and our Inception Capsule network achieves competitive results without requiring data augmentation. In contrast to CapsNet, we used a ReLU activation function before the squashing function in all layers, and employed three routing iterations between the Inception PrimaryCapsules and the VesselCaps. We found that using subnetworks to reconstruct vessel image patches as in [11] did not improve the performance and thus we did not use it. For vessel segmentation, we used one Capsule for the VesselCaps layer, and for centerline extraction (which includes segmentation) we used two Capsules.

**Fig. 3:**
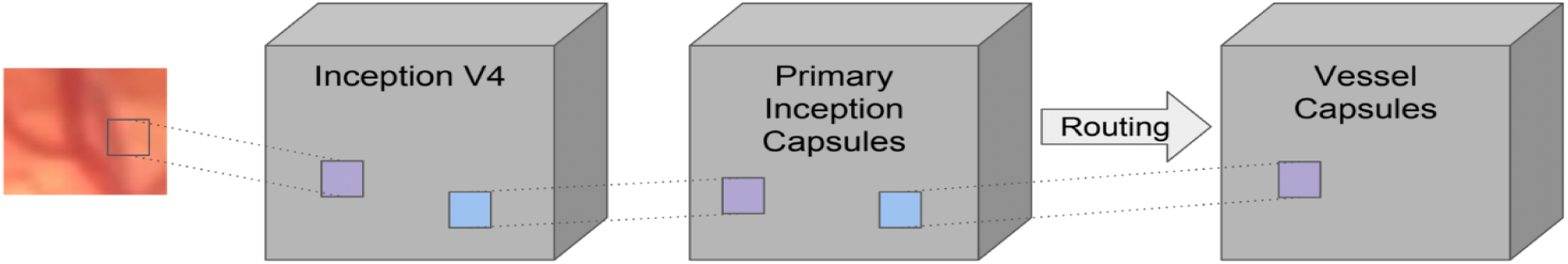
Proposed Inception Capsule architecture for retinal blood vessel segmentation and centerline extraction.

### 2.3. Training of the network

To train our Inception Capsule network, we extracted for each image 80.000 random image patches of size 27 × 27 pixels without resampling. Each image patch is entirely located in the field of view (FOV) of the retinal disk and was labeled corresponding to the class of its central pixel. For testing, all pixels in an image were classified. The intensities within the image patches were zero-centered by subtracting the mean and additionally scaled by the standard deviation. We trained our model without data augmentation since we found that this did not improve the performance. All experiments were conducted using ADAM optimization with a fixed learning rate of 0.0001. The model was trained for 65 epochs with a batch size of 32 by using the margin loss as described in [11].

## 3. EXPERIMENTAL RESULTS

We assessed the performance of our proposed Inception Capsule network based on the publicly available DRIVE dataset [14]. This dataset contains ground truth segmentation masks for retinal vessels as well as the FOV of the retinal disk, which were manually annotated by experts. The images have a size of 565 × 584 pixels. The dataset is split into 20 images for training and 20 images for testing. For centerline extraction, we applied skeletonization to the segmentation masks to determine the centerline ground truth. For both segmentation and centerline extraction, we applied 5-fold cross-validation on the training images and combined the test predictions of each fold in an ensemble. We evaluated our model using the following metrics: Accuracy (Acc), Area under the curve (AUC), Sensitivity (Sens), and Specificity (Spec). In Table 1 we provide the results of our method for vessel segmentation and a comparison with state-of-the-art methods. Fig. 4 shows an example result of our method. Our proposed Inception Capsule network yields an improvement compared to existing methods that classify central pixels ([4], [5], [10]) and achieves state-of-the-art results similar to deep autoencoders ([7], [8], [9]). For the Acc metric our method obtained the overall best result. Compared to CapsNet [11], the performance of our Inception Capsule network is significantly better. The results of our method for centerline extraction are given in Table 2 in comparison to a convolutional network similar to the one in [5] (Baseline), the Deep-FExt method in [6] that uses an Inception network, the U-Net [16], which is a deep autoencoder, and the CapsNet [11]. Our method significantly outperforms [5] and [11], and generally achieves better results than [6] and [16]. Computations were performed on an Intel Core i7-8700K CPU workstation with an Nvidia Titan X (Pascal). The implementation was done in Pytorch. The training took 3.75 days for 65 epochs and 32 minutes per test image.

**Table 1:** Performance of our method for vessel segmentation compared to state-of-the-art methods. Bold and underline represents the best result, and bold indicates the second best result.

| Method | Acc | AUC | Sens | Spec |
| --- | --- | --- | --- | --- |
| Melinščak [4] | 0.9466 | 0.9749 | - | - |
| Fu [5] | 0.9523 | - | 0.7603 | - |
| Li [7] | 0.9527 | 0.9738 | 0.7569 | 0.9816 |
| Dasgupta [8] | 0.9533 | 0.9744 | **0.7691** | 0.9801 |
| Yan [9] | **0.9542** | **0.9752** | 0.7653 | **0.9818** |
| Liskowski [10] | 0.9251 | 0.9738 | **0.9160** | 0.9241 |
| CapsNet [11] | 0.9292 | 0.9638 | 0.7614 | 0.9731 |
| **Proposed** | **0.9547** | **0.9750** | 0.7651 | **0.9818** |

**Table 2:** Performance of our method for vessel centerline extraction compared to state-of-the-art methods.

| Method | Acc | AUC | Sens | Spec |
| --- | --- | --- | --- | --- |
| Baseline [5] | 0.740 | 0.691 | 0.576 | 0.826 |
| Deep-FExt [6] | 0.877 | 0.845 | 0.710 | 0.823 |
| U-Net [16] | **0.891** | **0.878** | **0.741** | **0.848** |
| CapsNet [11] | 0.779 | 0.828 | 0.668 | 0.831 |
| **Proposed** | **0.907** | **0.882** | **0.752** | **0.842** |

**Fig. 4:**
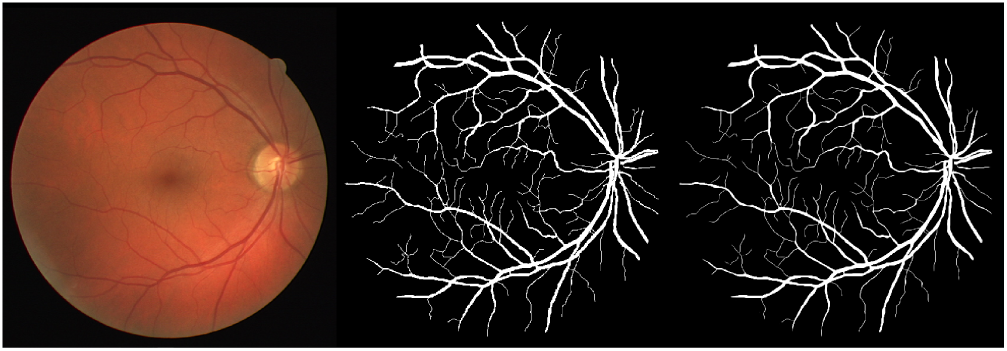
Segmentation result for an example image from the DRIVE dataset. (Left) retinal fundus image, (middle) manual ground truth, and (right) final segmentation result of the proposed method.

## 4. CONCLUSION

We have introduced a novel deep learning method for segmentation and centerline extraction of retinal blood vessels based on a combination of the Capsule network and the Inception architecture. An important property of our Inception Capsule network is its shallow architecture that has fewer parameters compared to state-of-the-art deep convolutional neural networks, while data augmentation is not required. A quantitative comparison with previous methods using the DRIVE dataset showed that our method yields competitive results for vessel segmentation and outperforms state-of-the-art methods for centerline extraction. In future work, we will investigate different extensions of our method to further reduce the computation time and we will apply our method to other datasets.

## REFERENCES

[1] S. J. Lee, C. A. McCarty, H. R. Taylor, and J. E. Keeffe, “Costs of mobile screening for diabetic retinopathy: A practical framework for rural populations,” Australian Journal of Rural Health, vol. 9, no. 4, pp. 186–192, 2001.

[2] N. Patton, A. Pattie, T. MacGillivray, T. Aslam, B. Dhillon, A. Gow, J. M Starr, L. J. Whalley, and I. Deary, “The association between retinal vascular network geometry and cognitive ability in an elderly population,” Investigative Ophthalmology & Visual Science, vol. 48, no. 5, pp. 1995–2000, 2007.

[3] K. He, X. Zhang, S. Ren, and J. Sun, “Deep residual learning for image recognition,” in Proc. CVPR 2016, pp. 770–778, IEEE, 2016.

[4] M. Melinščak, P. Prentašić, and S. Lončarić, “Retinal vessel segmentation using deep neural networks,” in Proc. VISAPP 2015, vol. 1, pp. 577–582, SciTePress, 2015.

[5] H. Fu, Y. Xu, S. Lin, D. W. K. Wong, and J. Liu, “Deep-vessel: Retinal vessel segmentation via deep learning and conditional random field,” in Proc. MICCAI 2016, pp. 132–139, Springer, 2016.

[6] G. Tetteh, M. Rempfler, C. Zimmer, and B. H. Menze, “Deep-FExt: Deep feature extraction for vessel segmentation and centerline prediction,” in Proc. MLMI 2017, pp. 344–352, Springer, 2017.

[7] Q. Li, B. Feng, L. Xie, P. Liang, H. Zhang, and T. Wang, “A cross-modality learning approach for vessel segmentation in retinal images.,” IEEE Trans. on Med. Imag., vol. 35, no. 1, pp. 109–118, 2016.

[8] A. Dasgupta and S. Singh, “A fully convolutional neural network based structured prediction approach towards the retinal vessel segmentation,” in Proc. ISBI 2017, pp. 248–251, IEEE, 2017.

[9] Z. Yan, X. Yang, and K. T. Cheng, “Joint segment-level and pixel-wise losses for deep learning based retinal vessel segmentation,” IEEE Trans. Biomed. Eng., vol. 65, no. 9, pp. 1912–1923, 2018.

[10] P. Liskowski and K. Krawiec, “Segmenting retinal blood vessels with deep neural networks,” IEEE Trans. on Med. Imag., vol. 35, no. 11, pp. 2369–2380, 2016.

[11] S. Sabour, N. Frosst, and G. E. Hinton, “Dynamic routing between capsules,” Advances in Neural Information Processing Systems, pp. 3856–3866, 2017.

[12] G. E. Hinton, S. Sabour, and N. Frosst, “Matrix capsules with EM routing,” in Proc. International Conf. on Learning Representations, pp. 3859–3869, 2018.

[13] C. Szegedy, W. Liu, Y. Jia, P. Sermanet, S. Reed, D. Anguelov, D. Erhan, V. Vanhoucke, and A. Rabinovich, “Going deeper with convolutions,” in Proc. CVPR 2015, pp. 1–9, IEEE, 2015.

[14] J. Staal, M. D. Abramoff, M. Niemeijer, M. A. Viergever, and B. van Ginneken, “Ridge-based vessel segmentation in color images of the retina,” IEEE Trans. on Med. Imag., vol. 23, no. 4, pp. 501–509, 2004.

[15] C. Szegedy, S. Ioffe, V. Vanhoucke, and A. A. Alemi, “Inception-v4, Inception-ResNet and the impact of residual connections on learning.,” in Proc. AAAI, vol. 4, pp. 4278–4284, AAAI Press, 2017.

[16] O. Ronneberger, P. Fischer, and T. Brox, “U-net: Convolutional networks for biomedical image segmentation,” in Proc. MICCAI 2015, pp. 234–241, Springer, 2015.

